# Lack of evidence for a role of anthrax toxin receptors as surface receptors for collagen VI and for its cleaved off C5 domain (endotrophin)

**DOI:** 10.1101/2021.10.29.466413

**Authors:** Matthias Przyklenk, Stefanie Elisabeth Heumüller, Steffen Lütke, Gerhard Sengle, Manuel Koch, Mats Paulsson, Alvise Schiavinato, Raimund Wagener

## Abstract

The widely expressed microfibril-forming collagen VI is subject to proteolytic cleavage and it has been proposed that the cleaved off C-terminal Kunitz domain (C5) of the α3 chain is an adipokine important for tumor progression and fibrosis. Under the name “endotrophin” the C5 fragment has also been shown to be a potent biomarker for fibro-inflammatory diseases. However, the biochemical mechanisms behind endotrophin activity have not been investigated. In earlier studies, the anthrax toxin receptor 1 was found to bind to C5, but this potential interaction has not been further studied. Given the proposed physiological role of endotrophin we aimed to determine how the endotrophin signal is transmitted to the recipient cells. Surprisingly, we could not detect any interaction between endotrophin and anthrax toxin receptor 1 or its close relative, anthrax toxin receptor 2. Moreover, we could not detect binding of fully assembled collagen VI to either anthrax toxin receptor. We also performed similar experiments with the collagen VI surface receptor NG2 (CSPG4). We could confirm that NG2 is a collagen VI receptor that binds to assembled collagen VI, but not to the cleaved C5/endotrophin. A cellular receptor for C5/endotrophin therefore still remains elusive.

## Introduction

Collagen VI is a microfibrillar protein expressed in most tissues and is there involved in anchoring large interstitial structures and cells (1). It is an unusual collagen in that it is predominantly composed of VWA domains and contains only a short collagenous domain. The VWA domains are crucial for collagen VI assembly (2) and for the interaction with other extracellular matrix proteins (3). The major form of collagen VI consists of α1, α2 and α3 chains, but in some cases the α3 chain is replaced by an α4, α5 or α6 chain (4). The α3-α6 chains are larger than the α1 and α2 chains and possess up to 10 tandem VWA domains in their N-terminal region.

The assembly of collagen VI microfibrils is a complex multi-step process. After the folding of heterotrimeric monomers (α1α2α3) these form antiparallel dimers (5) that finally assemble laterally to tetramers and are then secreted. Microfibrils form by intercalating end to end associations that are accompanied by proteolytic processing of a large C-terminal globular region of the α3 chain (6) which is absent in mature microfibrils (6, 7). This results in the release of the short most C-terminal C5 domain, a Kunitz domain, which was also termed endotrophin and proposed to be an adipokine (8, 9). Recently it was shown that endotrophin can be released by BMP1 (7) or MMP14 (10). However, the free endotrophin fragment is much less abundant than larger C5-containing fragments (7).

Collagen VI is linked to several diseases and, most prominently, mutations in *COL6A1, COL6A2* or *COL6A3* lead to myopathies (11). Bethlem myopathy and Ullrich congenital muscular dystrophy are the most frequent, inherited both in an autosomal dominant and recessive manner and represent, respectively, the mild and severe end of the disease spectrum (11). Moreover, collagen VI mutations can cause autosomal dominant limb-girdle muscular dystrophy (12) and autosomal recessive myosclerosis (13). Recessive mutations in *COL6A3* have been also linked to isolated dystonia (14).

The pathomechanisms leading to collagen VI related myopathies have been studied in mouse models. Mice lacking *Col6a1* display a phenotype reflecting Bethlem myopathy due to an abnormal opening of the mitochondrial pore and a disturbed autophagy in skeletal muscle (15, 16). However, it is still unclear how the lack of extracellular collagen VI leads to an intracellular phenotype. Integrins (17), the cell surface proteoglycan NG2 (18), and the anthrax toxin receptors 1 (19) and 2 (20) have been reported to be collagen VI receptors. Interestingly, the latter two have been linked to collagen VI in fibroproliferative diseases. Mutations in the anthrax toxin receptors cause GAPO syndrome (anthrax toxin receptor 1, ANTXR,1 also known as TEM8) (21) or infantile hyaline fibromatosis (anthrax toxin receptor 2, ANTXR2, also known as CMG2) (22) that are both accompanied by excessive deposition of collagen VI (23, 24). Primary fibroblasts isolated from *Antxr1* deficient mice display an up-regulated expression of collagen VI and collagen I pointing to the antiproliferative function of this receptor (24). It is however unclear if the loss of binding of collagen VI to Antxr1 interrupts a feedback loop. Moreover, even a conditional knock-out of *Antxr1* in vascular endothelial cells causes severe skin fibrosis and increased expression of collagen VI (24), indicating a complex cross-talk between tissues also involving VEGF signalling (24). The fibrotic condition could also be the consequence of a reduced degradation, as the activity of the matrix-degrading enzyme MMP2 is strongly decreased (24). Mice deficient in Antxr2 show a massive deposition of collagen VI in the uterus leading to an infertility which is rescued in *Antxr2/Col6a1* double deficient mice. In contrast to in *Antxr1* deficient mice, collagen VI accumulation is probably not caused by a reduced degradation by metalloproteinases in *Antxr2* deficient mice. However, there is evidence that Antxr2 is involved in endocytosis and lysosomal degradation of collagen VI (20). In contrast to Antxr1, which was shown to bind to the C5 domain of the α3 chain (19), it was proposed that Antxr2 binds to the triple helical domain of collagen VI (20).

For endotrophin, the cleaved-off C5 domain, growing evidence has been provided that point to important roles in tumour progression, fibrosis, inflammation and insulin resistance (8, 9). However, the cellular receptor/s responsible for mediating these effects are still not known. A proposed interaction between the Antxr1 and the C5 domain was originally based on a yeast two-hybrid screen and co-immunoprecipitation of overexpressed Antxr1 and C5, but was not studied in detail (19). Recently, serum levels of endotrophin have been shown to be valuable biomarkers for several pathological conditions (25). The combination of a proposed important physiological role of endotrophin and its growing importance as a biomarker led us to study the interaction with its proposed receptors in more detail. Moreover, the second anthrax toxin receptor, Antxr2, which shares a high homology with Antxr1, was shown to act as a receptor for collagen VI, even if it remains unknown which part of the collagenous domain is mediating such an interaction (20). Therefore, we revisited the interaction of the C5 domain (endotrophin) and of collagen VI with the anthrax toxin receptors and present data that substantially challenge current concepts.

## Results

### The VWA domains of the anthrax toxin receptors do not bind to the proteolytically released C5 domain of the collagen VI α3 chain

Since the cellular receptor/s mediating the biological activity of endotrophin — the cleaved off C5 domain of collagen VI — is/are not known, we revisited the well accepted binding of C5 to Antxr1 (19) by performing direct protein-protein interaction studies. Such studies, to our knowledge, have not been performed earlier. For that purpose, we expressed and purified recombinant murine C5 and the VWA domains of both Antxr1 and Antxr2 in HEK293T cells (Figure 1A). We decided to use the anthrax toxin receptor VWA domains as they are proposed to be the major extracellular ligand-binding regions of the two receptors and known to be sufficient to bind to the anthrax toxin (26, 27). In our analysis we included the VWA domain of Antxr2 even if it was proposed that collagen VI binds to this receptor via its collagenous domain (20). As a positive control we used the protective antigen (PA) subunit of the anthrax toxin, the pathogenic and well characterized ligand of the anthrax toxin receptors (28). By performing direct interaction studies using surface plasmon resonance, we confirmed that both VWA domains can bind to PA in a cation-dependent manner, with the VWA domain of Antxr2 showing a higher binding affinity than the same domain of Antxr1 (K_D_s: 6.45 ± 1.08 nM and 182 ± 64.3nM, respectively) as previously reported (29). Surprisingly, using the same assay under the same conditions, we could not detect any direct interaction between the VWA domain of Antxr1 and the C5 domain. Similarly, also the VWA domain of Antxr2 did not show any binding to C5 (Figure 1B).

**Figure 1.**
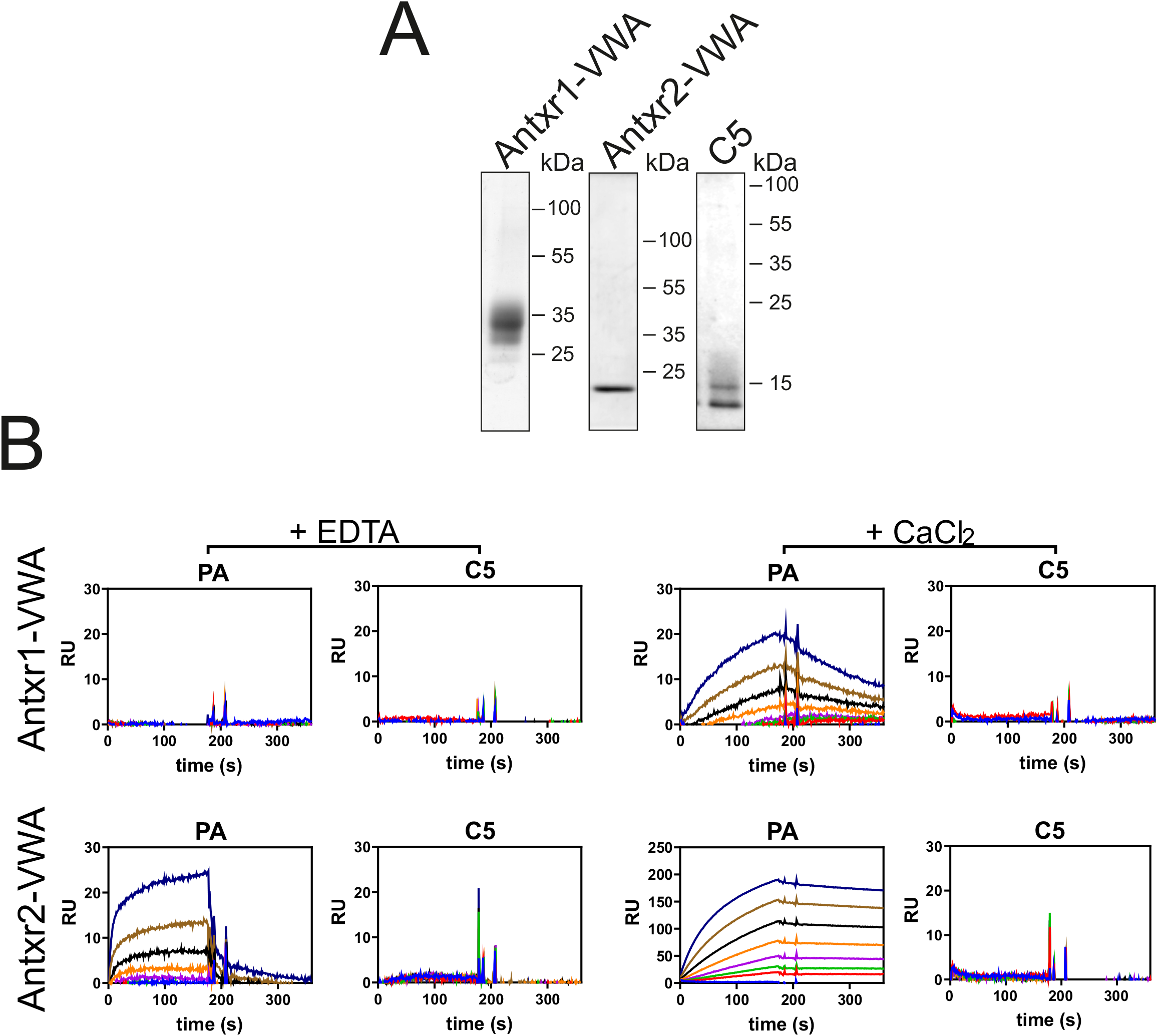
PA, but not the C5 domain of the collagen VI α3 chain binds to the VWA domains of anthrax toxin receptors. (**A**) Coomassie Blue stained SDS/polyacrylamide gel electrophoresis with the recombinant proteins used for the binding experiments. The upper C5 band is likely due to different glycosylation of the recombinant protein. (**B)** SPR sensorgrams obtained either in presence of EDTA or CaCl_2,_ of the interaction between murine Antxr1 or Antxr2 VWA-domains (immobilized on the sensor chips in similar quantities) and Protective Antigen (PA) or murine C5 (flown over as soluble ligands) in a dilution series from 0 to 320 nM. Ligand binding to the immobilized proteins on the chip is shown as response units on the y-axis (*n=3*).

### Plasma membrane located, full-length anthrax toxin receptors do not mediate cell binding to the recombinant C5 domain

The fact that we could not detect a direct binding of the C5 domain to the principal binding domain of the Antxr1 was surprising and in conflict with results obtained by others (19). Our contradictory findings made it necessary to unequivocally confirm this result by using alternative methods employing a biologically more relevant setting. Therefore, we generated HEK293T cell lines stably overexpressing full-length Antxr1 or −2 on the plasma membrane to test if they support cell binding to the C5 domain. To first validate the system, we treated the cells for 1 hour at 37° C with 5 μg/ml PA in serum-free medium. When using a PA-specific antibody, staining of control HEK293T cells that had been transfected with an empty vector, did not reveal any immunoreactivity. In contrast, when PA was added to Antxr1 or Antxr2 expressing cells we could detect a strong PA signal colocalizing with the two receptors on the cell surface (Figure 2A). However, when the same cells were treated with an equivalent concentration of the C5 domain and stained with a C5-specific antibody, we could not detect any positive signal on control cells or on cells overexpressing the two anthrax toxin receptors (Figure 2B). To further confirm that Antxr1 and Antxr2 act as surface receptors for PA but not for C5, we once more treated HEK293T cells with the two ligands and analysed the amount of cell-bound protein by immunoblotting. Consistent with our immunofluorescence results, we could recover an increased amount of PA from the cell lysate fractions upon Antxr1 or Antxr2 overexpression, while we found that the C5 domain was only present in the culture medium and not in the cell layer of the three cell lines, irrespective of the expression of the anthrax toxin receptors. Altogether, these results show that expression of the full-length anthrax toxin receptors mediates the binding of PA, but not of the C5 domain of the collagen VI α3 chain, to HEK293T cells. To exclude the possibility that C5 does not bind to the cell surface of HEK293T cells because of the lack of a co-receptor, we performed similar experiments using CHO-K1 cells, since it is known that these can bind and uptake the anthrax toxin (28). For this purpose, we generated CHO-K1 cell lines overexpressing full-length Antxr1 and Antxr2 receptors and found that, while the overexpression of the two receptors greatly increased the capacity of PA to bind to the cells, they were completely unable to mediate the binding of C5 to the cell surface (Supplementary figure 1). Importantly, using the same assays (immunostaining or immunoblot) we could not detect any interaction of the recombinant C5 domain with the cell surface of mouse primary dermal fibroblasts, MC3T3-E1, RPE-1, U2OS, MCF-7 or MDA-231-MB cells (not shown).

**Figure 2.**
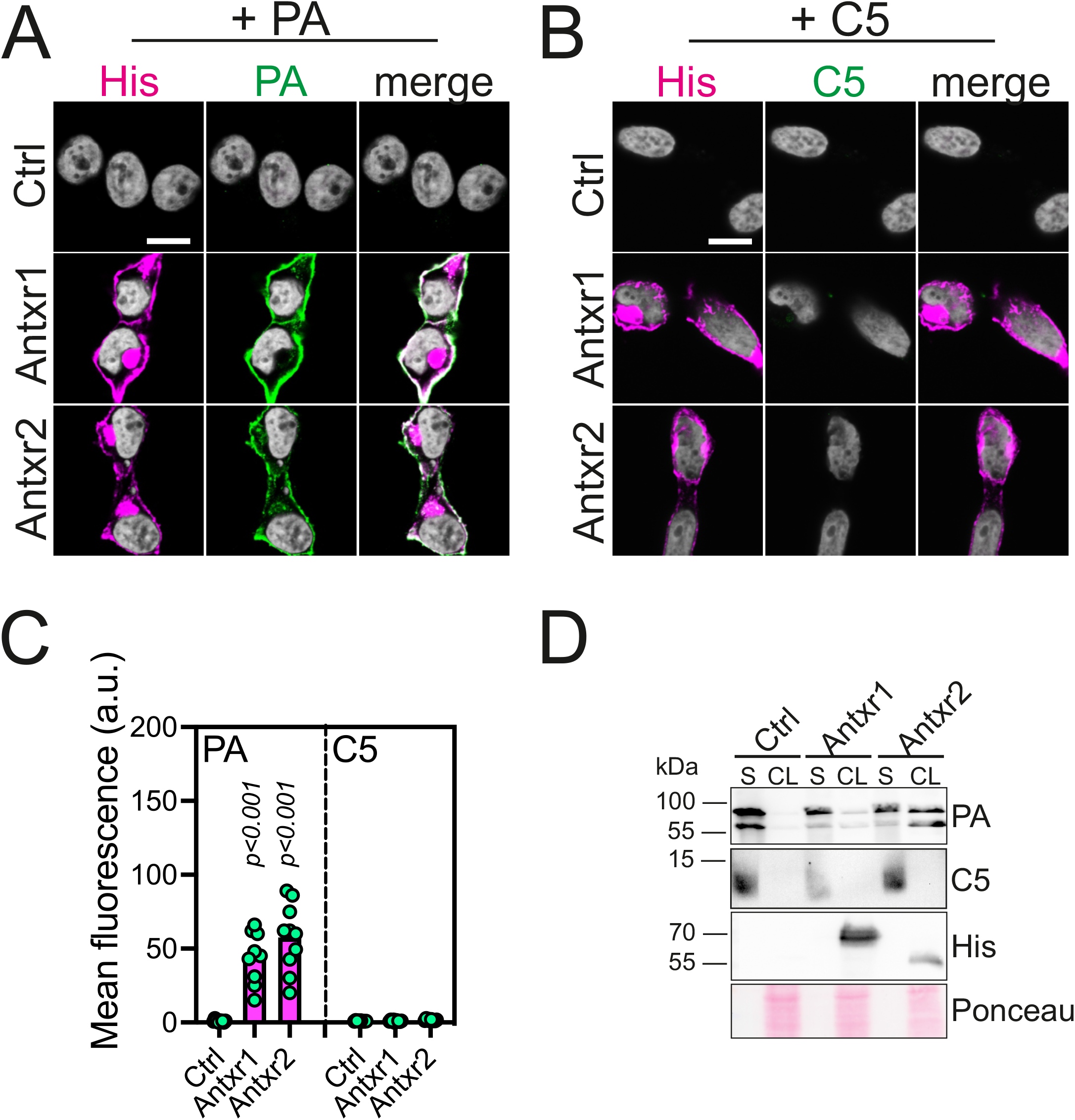
Anthrax toxin receptors support the binding of PA, but not of C5 to the surface of HEK293T cells. (**A-B)** Confocal microscopy pictures of control and His-tagged Antxr1 or Antxr2 overexpressing HEK293T cells labelled with the indicated antibodies. Cells were treated with 5 μg/ml PA (A) or human C5 (B) in serum-free medium for 1 h at 37° C, washed and then fixed. Similar experiments were also carried out using recombinant murine C5 with identical results (not shown). **(C)** Quantification of the mean fluorescence intensity of the immunolabeled cells as in A and B. **(D)** Control and anthrax toxin receptors overexpressing HEK293T cells were treated with 1 μg/ml PA or murine C5 for 1 h at 37° C and then washed with PBS before collecting the culture supernatants (S) and harvesting the cell layers (CL). Samples were immunoblotted using the indicated antibodies. Each experiment was performed at least three times with similar results. Scale bars: 10 μm

### The cell surface proteoglycan NG2, and not the anthrax toxin receptors, supports cell binding to collagen VI, but not to the isolated C5 domain

Our previous experiments did not provide evidence supporting a role of anthrax toxin receptors as cell-surface receptors for the recombinant C-terminal C5 domain (endotrophin) of the collagen VI α3 chain. We therefore performed further experiments with a more physiological source of C5. For this purpose, we collected the conditioned medium of MC3T3-E1 osteoblasts grown in the presence of ascorbate, since we have previously found that these cells secrete collagen VI tetramers and form an abundant collagen VI microfibrillar network (30). Next, we treated control and anthrax toxin receptor overexpressing HEK293T cells with the conditioned medium and stained them with the C5 antibody. As a positive control, we also generated HEK293T cells stably expressing the integral plasma membrane proteoglycan NG2 (CSPG4), an established collagen VI receptor (31). C5-containing collagen VI molecules present in the osteoblast medium bound scarcely to control HEK293T cells and, similar to what we observed using the recombinant C5 domain, overexpression of the two anthrax toxin receptors did not increase the interaction of collagen VI with the cells. In contrast, using the same antibody, we detected an intense signal on the cell surface of NG2 expressing cells (Figure 3A-B). The C5 domain is progressively cleaved off from the α3 chain after the secretion of collagen VI tetramers (7, 32). Therefore, the signal detected on the NG2-expressing cells could represent either the released C5 domain, C5-containing collagen VI molecules or fragments thereof. To distinguish between these possibilities, we stained the same cells with an antibody (α3N) specific for the N-terminal region of the α3 chain that is not cleaved from the rest of the molecule. Using this antibody, we observed no or very scarce reactivity on control cells and on cells expressing the anthrax toxin receptors, while NG2-expressing cells were intensively labelled with a signal that colocalized completely with that obtained with an antibody against C5 (Figure 3A-B). Another co-staining experiment with the different cell lines, using the α3N antibody together with an antibody specific for the α1 chain of collagen VI (α1C), gave similar results (Supplementary Figure 2). Next, we analysed extracts of the cells incubated with the conditioned medium by immunoblotting. Using an antibody specific for the α1 chain we found that only NG2-expressing cells showed an increased binding. Strikingly, using the C5 antibody on the same samples, we could confirm that the cells expressing NG2, but not Antxr1 or Antxr2, showed enhanced binding to the fully assembled C5-containing collagen VI tetramers, while the soluble, cleaved off C5 domain did not bind to any of the cell lines (Figure 3C). Since these results are in disagreement with previous studies proposing a role for Antxr2 as the collagen VI receptor, we sought to further verify our findings. Toward this aim we performed pull-down assays using full-length anthrax toxin receptors and conditioned medium of MC3T3-E1 cultures. However, despite the fact that we performed the assay multiple times under different conditions, we could not pull-down collagen VI or the cleaved off C5 domain with the anthrax toxin receptors, while PA was consistently pulled down (Figure 3D). On the other hand, we could co-precipitate collagen VI from MC3T3-E1 conditioned medium in a similar assay by using the recombinant ectodomain of NG2 (33) (Figure 3E). Our results indicate that collagen VI tetramers bind only scarcely to HEK293T cells, and that NG2, but neither Antxr1 nor −2, is able to endow the cells with an increased binding activity. It has been previously shown that NG2 is able to interact with both pepsin-solubilized collagen VI, lacking its globular, non-collagenous domains, and the recombinant collagen VI α2 chain while the N-terminal region of the α3 chain showed no binding (34). However, it is not known if the C-terminal part of the α3 chain may also participate in the interaction. Therefore, to determine if the isolated C5 domain could be a ligand for NG2, and use this receptor to convey signals into the cell, we treated in parallel NG2-expressing cells with either recombinant C5 domain or the conditioned medium from osteoblast cultures or conditioned medium from collagen VI α2 deficient osteoblasts and stained them with the C5 antibody. We found that while cells treated with the conditioned medium showed a strong signal on the cell surface, no staining was present on the cells treated with the recombinant C5 protein, showing that the C5 domain alone does not possess the ability to bind to NG2. Moreover, conditioned medium derived from the α2 deficient osteoblast cultures that contain only single α1 and α3 chains of collagen VI, but no assembled tetramers, showed no significant binding of collagen VI molecules to the cell surface, implying that the α2 chain or a full tetramer conformation is needed to allow efficient binding to NG2. (Figure 3F-G).

**Figure 3.**
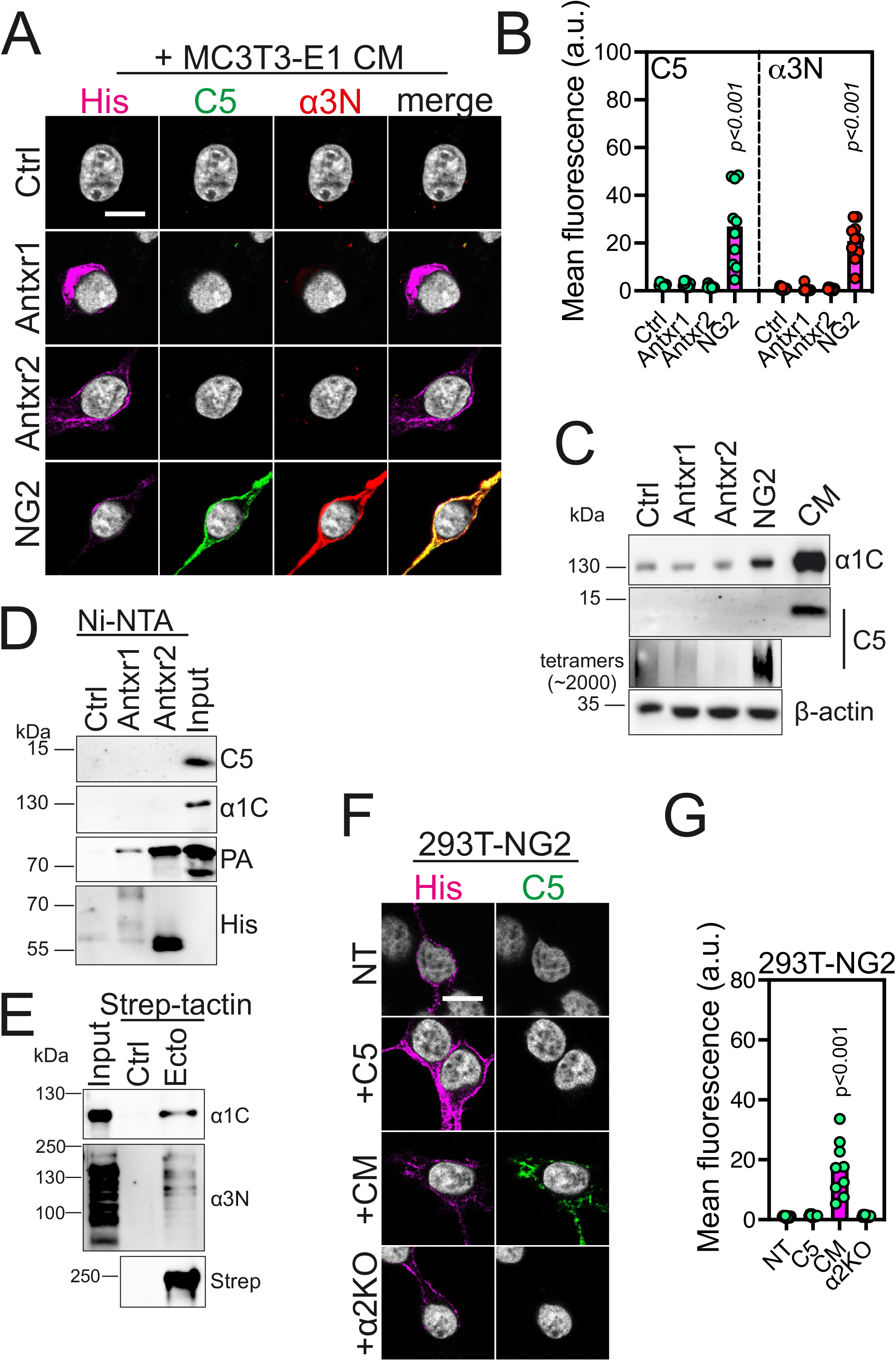
NG2, but not anthrax toxin receptors, mediates the binding of collagen VI to the cell surface. (**A)** Confocal microscopy pictures of control, His-tagged Antxr1, Antxr2 or NG2 overexpressing HEK293T cells that were incubated for 1 hour at 37° C with collagen VI-containing serum-free conditioned medium derived from MC3T3-E1 osteoblasts. After washing, cells were fixed and immunolabeled with the indicated antibodies. **(B)** Quantification of the mean fluorescence intensity of the immunolabeled cells as in A. **(C)** Control HEK293T cells and cells expressing the indicated receptors were treated with the conditioned medium as in A. After washing, cell lysates were harvested and analysed by immunoblotting using the indicated antibodies to detect collagen VI moieties bound to the cell layers. For detection of the α1 chain (top panel) by immunoblot, samples were resolved on a 10 % acrylamide gel under reducing conditions; for the cleaved off C5 domain (middle panel) a 15 % acrylamide gel was used under non-reducing conditions. For immunoblotting of collagen VI tetramers (bottom panel), samples were submitted to composite agarose/acrylamide gel electrophoresis under non-reducing conditions and immunoblotted with the C5 antibody. β-actin was used as a loading control. **(D)** Cell lysates of control HEK293T cells and of cells expressing His-tagged Antxr1 or Antxr2 were incubated with Ni-NTA agarose beads and with either collagen VI containing conditioned medium or PA spiked into the same medium. After washing the beads, bound proteins were eluted in SDS-containing sample buffer and analysed by immunoblotting with the indicated antibodies. **(E)** Strep-Tactin coupled sepharose beads were incubated with recombinant strep-tagged NG2-ectodomain (Ecto) or without any protein (Ctrl). The beads were afterwards incubated with collagen VI containing conditioned medium. After washing, the proteins bound to the beads were eluted in SDS-containing sample buffer and immunoblotted with the indicated antibodies **(F)** Confocal microscopy pictures of NG2-expressing HEK293T cells incubated for 1 hour with either recombinant C5 or collagen VI-rich conditioned medium derived from MC3T3-E1 osteoblasts, or conditioned medium derived from collagen VI α2-chain deficient MC3T3-E1 osteoblast cultures. After washing, cells were labelled with the indicated antibodies. **(G)** Quantification of the mean fluorescence intensity of labelled cells as in E. CM: conditioned medium; NT: not treated. Scale bars: 10 μm

**Figure 4.**
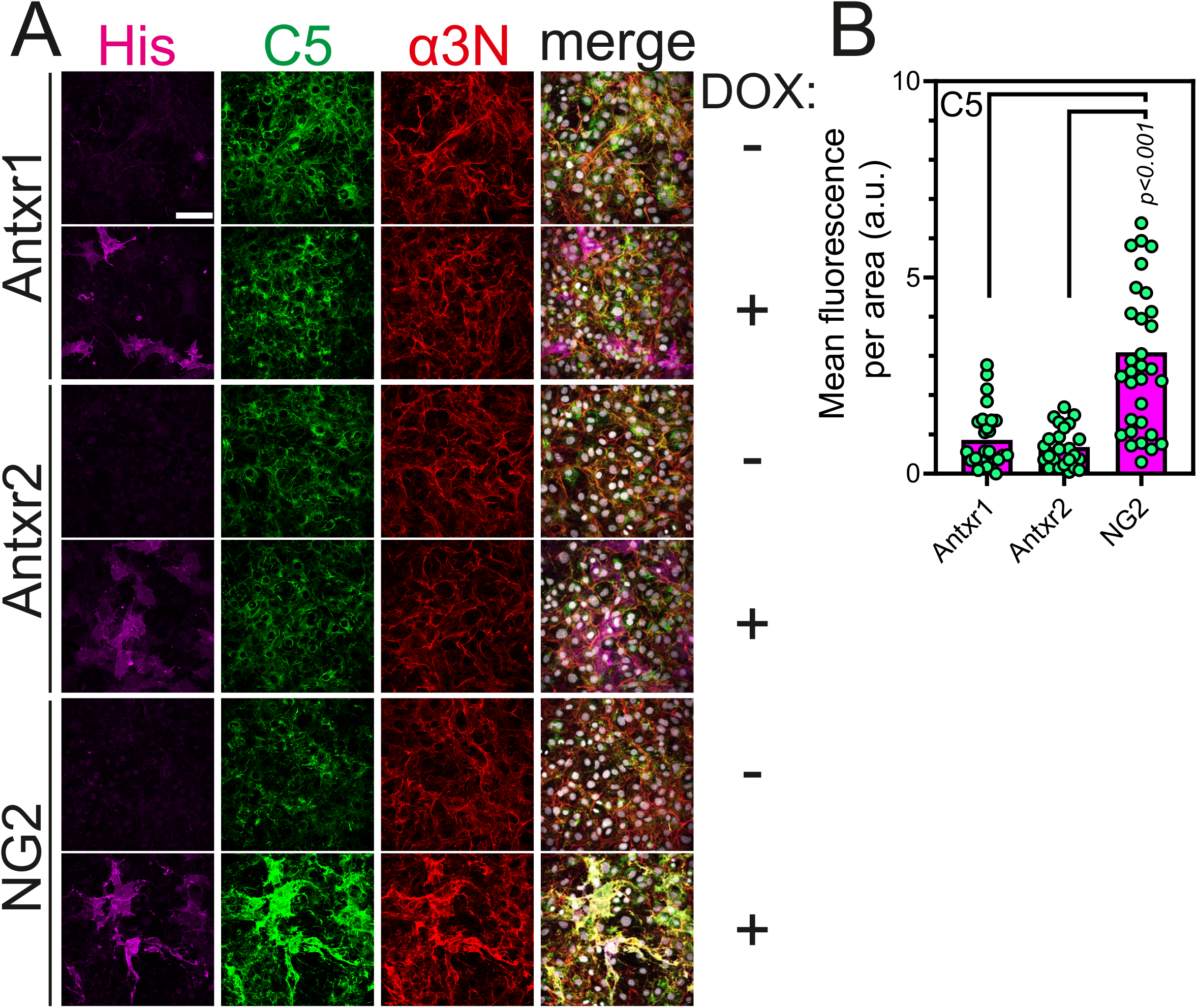
Expression of NG2, but not of anthrax toxin receptors, causes retention of type VI collagen on the surface of MC3T3-E1 cells. **(A)** Confocal microscopy images of His-tagged Antxr1, Antxr2 or NG2 overexpressing MC3T3-E1 cells labelled with the indicated antibodies. Cells were grown for 5 days in the presence of ascorbate with or without doxycycline to induce the expression of the receptors. **(B)** Quantification of the mean C5 fluorescence intensity of labelled cells as in A. Mean fluorescence was measured in receptor expressing cells and related to area size. Scale bar: 100 μM

Our results so far have failed to demonstrate a role of anthrax toxin receptors as surface receptors for collagen VI or for C5. Next, we sought to investigate if anthrax receptors may interfere with collagen VI production in a cell-autonomous manner when expressed in collagen VI producing cells. Therefore, we generated MC3T3-E1 osteoblasts with inducible expression of His-tagged full-length Antxr1 and −2. Again, we included NG2 in our experiments since it has been previously shown that expression of NG2 in collagen VI producing cells caused retention of collagen VI on the cell surface (18). Cells were grown in the presence of ascorbate for 5 days with or without doxycycline and then analyzed by immunofluorescence. Upon doxycycline treatment expression of the three receptors was strongly induced in MC3T3-E1 cells as shown by staining with an antibody against the His-tag. To study the effect on collagen VI deposition we stained the cells with the C5 and α3N antibodies. The α3N antibody mainly stained extracellular collagen VI fibers incorporated into the ECM of the MC3T3-E1 osteoblasts, while the C5 antibody gave a more prominent intracellular signal, as previously described also in other cell types (35). Upon expression of the anthrax toxin receptors, we could not detect any significant differences in collagen VI localization and deposition. Moreover, no colocalization between anthrax toxin receptors and C5 or α3N was evident. On the other hand, NG2 expression caused a marked retention of collagen VI on the cell surface, where the C5 and α3N signals were largely overlapping with the signal for NG2.

## Discussion

C5, also named endotrophin, the C-terminal cleavage product of the collagen VI α3 chain has received a great deal of attention as it was proposed to act as an adipokine that is involved in tumorigenesis and fibrogenesis. Moreover, endotrophin has been shown to be a valuable biomarker for fibrotic conditions (for review see (36)), in this context also named PRO-C6. Long before the C5 domain was recognized as an adipokine and biomarker, it was shown that it binds to the Antxr1(19). To date no other receptor for endotrophin has been identified (36) and it is remarkable that although the physiological function of endotrophin has been extensively studied the interaction between endotrophin and a receptor that transmits the signal into the cell has not been further shown. The still enigmatic function of endotrophin is in part reminiscent of that of the adipokine adiponectin, where more than 25 years after its discovery (37) it is still not clear how and why it exerts multiple beneficial effects on various tissues and organs (38). However, in contrast to endotrophin adiponectin-binding partners, such as AdipoR1/2 (39) or T-cadherin (40), have been identified.

Therefore, after we showed that endotrophin (C5) is released by BMP1 (7) we carefully searched for receptor ligand interactions to gain better insight into the mechanism behind the proposed physiological responses to endotrophin. However, despite extensive attempts we were not able to find any evidence for a receptor interaction, although we applied not only methods to detect direct binding of recombinant proteins, but also performed cell culture-based interaction studies. It remains uncertain how the earlier results indicating a binding of the C5 domain to the Antxr1 were obtained. The understanding and reproduction of the former experiments is hampered by the fact that the exact borders of the C5 construct were not given by Nanda et al. (19). A 210 bp cDNA clone encoding the C-terminal end of the collagen VI α3 chain is mentioned. If this is consistent with a fragment encompassing the last 70 amino acid residues of the α3 chain, this fragment would start with E_3108_TDI. It would be slightly shorter than endotrophin released by BMP1 which starts with T_3101_EPL or endotrophin released by MMP14 which starts at A_3087_ RSA. Nevertheless, also the C5 fragment used by Nanda et al. most likely contained all six cysteine residues that are required for proper folding of a Kunitz domain (41). However, the yeast-two-hybrid screening that was used for the identification of the interaction may not be suited to study protein-protein interactions that naturally occur in the extracellular space. Structurally important posttranslational modifications of extracellular proteins, in particular the formation of disulphide bonds, are hindered by the reducing conditions in the nucleus where the interaction in yeast-two-hybrid assays actually takes place. The clear impact of reduction on the C5 structure is evident by the strong difference in its mobility on SDS-PAGE (35). Finally, the immunoprecipitation experiments that were performed to verify the yeast-two-hybrid result are lacking controls and are therefore of uncertain validity.

The fact that we did not observe binding of the C5 domain to a variety of cultured cells makes it difficult to understand the mechanism behind its proposed function as an adipokine. In particular, we could not see a binding or uptake of endotrophin by MCF-7 or MDA-MB-231 cells. This is remarkable as these cell lines have been shown to display an up-regulation of the epithelial mesenchymal transition markers Twist and Snail, and an enhanced cell migration upon endotrophin exposure (42). In line with our negative results, there are conflicting data on if collagen VI affects MDA-MB-231 cell migration not through endotrophin, but via interaction of the triple helical part of collagen VI with NG2 (43).

Nevertheless, when we used NG2 as a positive control we could confirm that NG2 binds to collagen VI, but also show that NG2 does not bind to endotrophin. This differs to our results on the proposed role of the Antxr2 in the uptake of collagen VI. Although it is evident that this receptor, like Antxr1, does not bind to endotrophin, it was surprising that we also could not detect binding of Antxr2 to collagen VI while it was proposed that Antxr2 is involved in the endocytosis and lysosomal degradation of assembled collagen VI tetramers (20). We could not provide evidence for a direct interaction between Antxr2 and collagen VI. If Antxr2, as proposed, is involved in the endocytosis of collagen VI it must contribute indirectly, perhaps by activation of proteases located at the plasma membrane (44).

In summary, we could not obtain evidence for binding of endotrophin to cells, which would be a prerequisite for its manifold proposed roles. It remains difficult to explain how endotrophin influences lipogenesis, lipolysis and inflammation (45), how it induces the transcription of pro-fibrotic and pro-inflammatory genes (9) or how it acts as a chemokine augmenting tumor growth (8). Our results, however, are consistent with the proposed role of endotrophin during the complex assembly of collagen VI (32, 46). It is therefore likely that endotrophin as a biomarker senses newly expressed collagen VI, which is a hallmark of fibrotic conditions.

## Experimental Procedures

### Antibodies

Affinity purified polyclonal antibodies against the C5 domain of the α3 chain, the C-terminal region of the α1 chain and the N-terminal region of the α3 chain of collagen VI were previously described (4, 7). The goat antibody against the anthrax protective antigen (PA) was from List Biological Laboratories and the mouse monoclonal antibody against the His-tag from Qiagen.

### Plasmids and recombinant proteins

Human and murine C5 domains were cloned as previously described (7) using the following primers: hC5: AAAGCTAGCGACAGAACCATTGGCTCTC(forward) and TTTGGATCCGGTTCCCATCACACTGAT(reverse); mC5: AAAGCTAGCAACAGAACCATTGTTTCT(forward) and TTTGGATCCAACTGTTAACTCAGGACTACACA(reverse). Both human and murine C5 sequences were chosen to start at the BMP1 cleavage site (7). Murine Antxr1-VWA (Gly34-Ile221) and Antxr2-VWA (Glu35-Ile221) were cloned using the following primers. Antxr1-VWA: AAAGCTAGCAGCTTGCTACGGAGGATTC(forward) and TTTGGATCCTTCGATGCAGGATTTCTT(reverse); Antxr2-VWA: AAAGCTAGCATCTTGCAAAAAAGCCTTC(forward) and TTTGGATCC TTCAGTACATGATTGAGC(reverse). Amplified PCR products were inserted into a modified pCEP-pu vector containing an N-terminal BM40 signal peptide as well as a C-terminal Twin-Strep-tag (47). Full-length murine Antxr1 and 2 were cloned using the following primers. Antxr1: CAATGCTAGCGGGCCGCCGCGAGGATG (forward) and CAATGGATCCGACAGAAGGCCTTGGAGGAGG (reverse); Antxr2: CAATGCTAGCCCAGGAGCAGCCCTC (forward) and CAATCTCGAGTTGATGTGGAACCCGGGAG (reverse). Amplified PCR products were inserted into a modified sleeping beauty vector (48). A construct coding for the full-length human NG2 cloned into the pEF6/MycHis-B vector was previously described (49). The NG2-ectodomain (33) was subcloned into the same modified sleeping beauty vector from the full-length construct using the following primers: GATGCTAGCGGCTTCCTTCTTCGGTGAGAAC(forward) and GAATGCGGCCGCGAACATGTTGGCCTCAAGG(reverse) Human and murine collagen VI C5 domains, anthrax toxin receptor VWAdomains and the NG2ectodomain were expressed and purified as previously described (7). Briefly, plasmids were introduced into HEK293T cells using the FuGENE 6 transfection reagent (Roche Applied Science). Cells were selected with puromycin (1 μg/ml), and the recombinant proteins were purified directly from serum-free culture medium. After filtration and centrifugation (1 h, 10,000 × *g*), the supernatants were applied to a Strep-Tactin column (1.5 ml; IBA GmbH) and eluted with 2.5 mM desthiobiotin, 10 mM Tris-HCl, pH 8.0. Protein purity and concentration was estimated via SDS-polyacrylamide gel electrophoresis followed by Coomassie blue staining.

### Cell culture

All cell lines used in this study were obtained from the American Type Culture Collection (ATCC). Cells were cultured in Dulbecco’s Modified Eagle’s Medium (DMEM) or Ham’s F-12 nutritional mix. MC3T3-E1 cells were cultured with Minimum Essential Medium (MEM) growth medium. All media were supplemented with 10 % FBS and 1 % penicillin/streptomycin. MC3T3-E1 cells lacking the α2 chain of collagen VI were previously described (30).

To generate stable cell lines, HEK293T and CHO-K1 cells were transfected with the FuGENE6 transfection reagent (Roche Applied Bioscience) and selected by the addition of puromycin (1 μg/ml and 10 μg/ml respectively) for full-length anthrax toxin receptors 1 and 2, or blasticidin for NG2. Collagen VI-rich conditioned medium was obtained by letting MC3T3-E1 cells grow to confluence in 100 mm culture dishes and afterwards shifting them to serum-free medium. Ascorbate (0.625 M L-ascorbate/1.125 M L-ascorbate-2-phosphate) was supplied every second day for a total of 7 days. Harvested conditioned media were briefly centrifuged at 13,000 x g to pellet dead cells and cell debris.

### Surface plasmon resonance (SPR) interaction assays

SPR experiments were performed as described using a BIAcore 2000 system (BIAcore AB, Uppsala, Sweden) (50). Recombinant VWA domains of Antxr1 and-2 were immobilized at 1000 reference units (RUs) to a CM5 sensor chip using the amine coupling kit following the manufacturer’s instructions (Cytiva). Interaction studies were performed by injecting 0-320 nM analyte in HBS-P buffer (0.01 M HEPES, pH 7.4, 0.15 M NaCl, 0.005% (v/v) surfactant P20) supplemented with 10 mM CaCl_2_ and HBS-EP (0.01 M HEPES, pH 7.4, 0.15 M NaCl, 3 mM EDTA, 0.005% (v/v) surfactant P20) (Cytiva). Kinetic constants were calculated by nonlinear fitting (1:1 interaction model with mass transfer) to the association and dissociation curves (BIAevaluation version 3.0 software). Apparent equilibrium dissociation constants (K_D_ values) were then calculated as the ratio of the dissociation rate constant (kd) and the association rate constant (ka). Experiments in the calcium-containing buffer were performed as triplicates.

### Binding assays with C5 domain, PA and collagen VI-rich conditioned medium

For immunofluorescence staining 6 × 10^4^ cells were plated on 12 mm glass cover slips in 24-well plates; for western blot analysis 1.5 × 10^5^ cells were plated in 12-well plates. Cells were left to grow for 24 hours before shifting them to serum-free conditions for another 24 hours. Cells were then incubated with recombinant proteins in serum-free medium (5 μg/ml for immunofluorescence or 1 μg/ml for immunoblotting) or with conditioned media for 1 h at 37° C. Afterwards, cells were washed twice with serum-free medium and fixed with 4% paraformaldehyde or lysed with RIPA buffer.

### Immunofluorescence microscopy

Fixed cells were permeabilized with 0.5% NP-40 for 10 min and blocked with 1 % FBS in PBS for 30 min. Cells were then incubated with primary antibodies diluted in 1 % FBS in PBS for 1 hour at RT, washed 3 x with PBS for 10 minutes, and incubated with appropriate highly cross-adsorbed secondary antibodies conjugated to Alexa Fluor 488, 555 or 647 (Thermo Fisher Scientific) and DAPI (0.1 μg/ml; Sigma-Aldrich) for 1 hour at RT. Cover slips were mounted with Fluorescence Mounting Medium (DAKO) on glass slides and images taken with a Leica TCS SP5 confocal microscope. For statistical evaluation, at least 10 images containing a single cell were randomly acquired and fluorescence intensities determined with the Leica LAS AF Lite software.

### Immunoblotting and pull-down assays

Cell lysates and cell culture supernatants were separated by 10 % or 15 % SDS-polyacrylamide gel electrophoresis under reducing conditions (non-reducing when using the antibody against C5), transferred to a nitrocellulose membrane and blocked with 5 % non-fat dry milk in 0.1 % Tween 20/TBS solution. The membranes were probed with primary goat antibodies against PA (1:5000, List Biological Laboratories), rabbit antibodies against C5 (1:1000, in-house generated), mouse antibodies against the His-tag (1:1000, Qiagen), rabbit antibodies against the C-terminal region of the collagen VI α1 chain (1:1000, in-house generated), guinea pig antibodies against the N-terminal region of the collagen VI α3 chain (1:500, in-house generated), or mouse antibodies against actin (1:5000, Millipore) and species corresponding horseradish peroxidase conjugated polyclonal secondary antibodies (DAKO) in blocking solution. Signals were detected by chemiluminescence (Cytivia, AmershamTM ECLTM prime western blot detection reagent). For immunoblotting of collagen VI tetramers, samples were subjected to electrophoresis on 0.5 % (w/v) agarose/2.4 % (w/v) polyacrylamide composite gels under non reducing conditions (30). For Anthrax receptor pulldown experiments, Indigo-Ni Agarose beads were used(Cube Biotech GmbH). Beads were washed and equilibrated in 0.15 M NaCl, 0.025 M Tris-HCl pH 7.5 prior to incubation with cell lysates from anthrax toxin receptor overexpressing cells (0.15 M NaCl, 0.025 M TrisHCl pH 7.5, 10% Glycerol, 1% NP-40, 1mM EDTA) for 3-6 hours. Afterwards the beads were washed three times (equilibration buffer with 0.1 % NP-40 and 0.02 M imidazol) and incubated with MC3T3-E1 conditioned media or conditioned media spiked with 1μg/ml PA overnight at 4° C. Elution of the bound material was performed after extensive washing by boiling the samples at 95° C with 2X Laemmli buffer. NG2 pulldown experiments were performed by binding recombinant NG2-ectodomain to Strep-Tactin coupled sepharose beads (IBA GmbH) for 3-6 hours. Afterwards the beads were incubated with collagen VI rich conditioned medium overnight. After extensive washing, bead bound material was eluted by boiling the samples at 95° C with 2x Laemmli buffer.

### Statistical analysis

GraphPad was used for generation of graphs and statistical analysis. For data comparison between multiple groups, one way ANOVA with Dunnett multiple testing correction was used, where a P value of ≤ 0.05 was considered as significant.

## Acknowledgment

We would like to thank Birgit Kobbe for her invaluable technical assistance. This work was supported by the Deutsche Forschungsgemeinschaft through project ID 384170921: FOR2722/B1-407164210 (MP and RW), FOR2722/B2 (MK) FOR2722/M2 and FOR2722/C2 (GS),

---

**Supplementary Figure 1.**
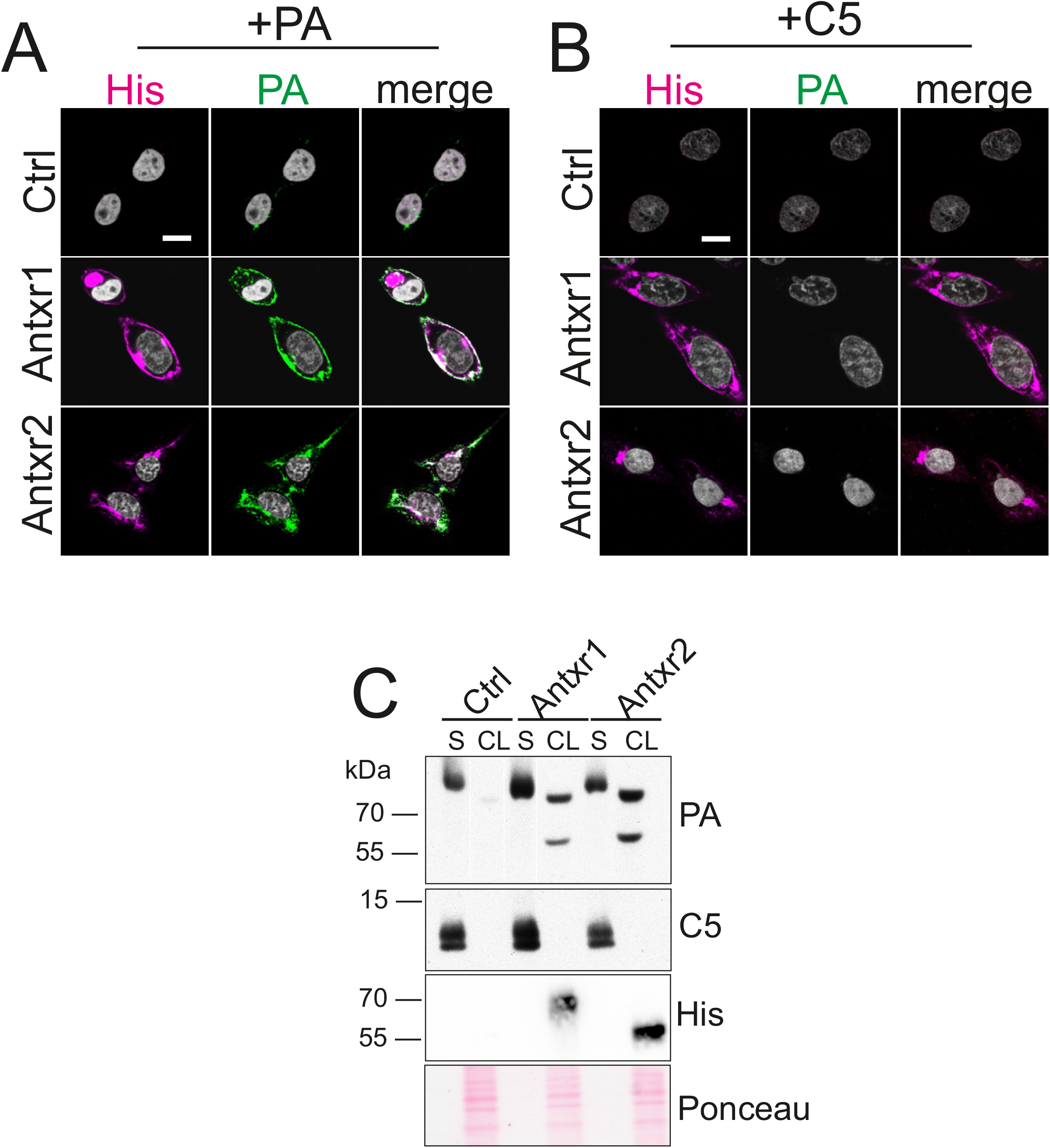
Anthrax toxin receptors support the binding of PA, but not of C5 to the surface of CHO-K1 cells. (**A-B)** Confocal microscopy images of control and His-tagged Antxr1 or Antxr2 overexpressing CHO-K1 cells labelled with the indicated antibodies. Cells were treated with 5 μg/ml PA (A) or human C5 (B) in serum-free medium for 1 h at 37° C, washed and then fixed. Similar experiments were also carried out using recombinant murine C5 with identical results (not shown). **(C)** Control and anthrax toxin receptors overexpressing CHO-K1 cells were treated with 1 μg/ml PA or human C5 for 1 h at 37° C and then washed with PBS before collecting the culture supernatants (S) and harvesting the cell layers (CL). Samples were immunoblotted using the indicated antibodies. Each experiment was performed two times with similar results. Scale bars: 10 μm

**Supplementary Figure 2.**
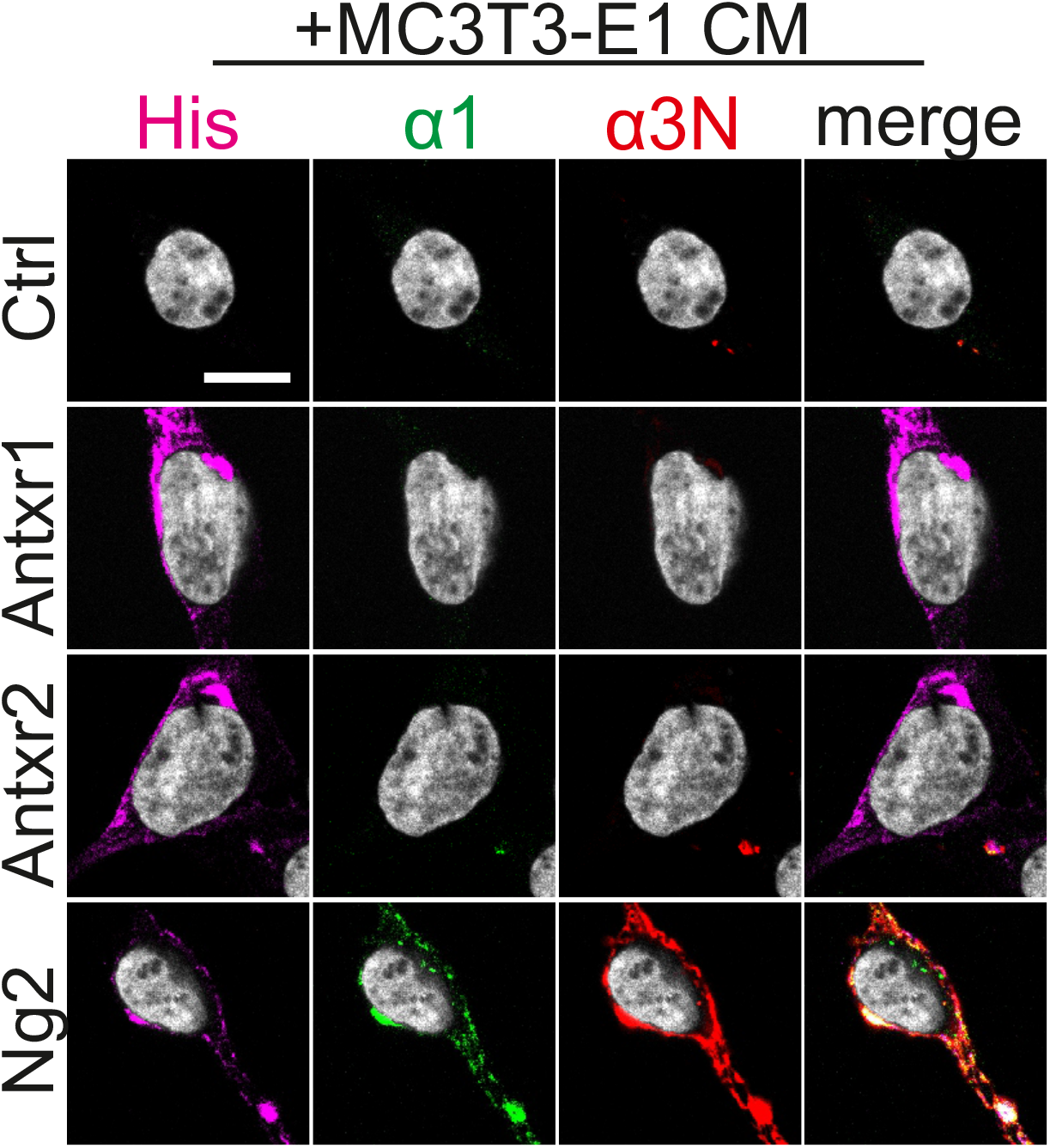
NG2, but not anthrax toxin receptors, mediates the binding of collagen VI to the cell surface. Confocal microscopy pictures of control and His-tagged Antxr1, Antxr2 or NG2 overexpressing HEK293T cells that were incubated for 1 h at 37° C with collagen VI-containing serum-free conditioned medium derived from MC3T3-E1 osteoblasts. After washing, cells were fixed and immunolabeled with the indicated antibodies. Scale bars: 10 μm

## References

1. Cescon M, Gattazzo F, Chen P, Bonaldo P. Collagen VI at a glance. Journal of Cell Science. 2015;128(19):3525–31.

2. Lamandé SR, Sigalas E, Pan TC, Chu ML, Dziadek M, Timpl R, et al. The role of the α3(VI) chain in collagen VI assembly. Expression of an α3(VI) chain lacking N-terminal modules N10-N7 restores collagen VI assembly, secretion, and matrix deposition in an α3(VI)-deficient cell line. Journal of Biological Chemistry. 1998;273(13):7423–30.

3. Wiberg C, Klatt AR, Wagener R, Paulsson M, Bateman JF, Heinegard D, et al. Complexes of matrilin-1 and biglycan or decorin connect collagen VI microfibrils to both collagen II and aggrecan. J Biol Chem. 2003;278(39):37698–704.

4. Maass T, Bayley CP, Mörgelin M, Lettmann S, Bonaldo P, Paulsson M, et al. Heterogeneity of Collagen VI Microfibrils: Structural Analysis of Non-collagenous Regions. J Biol Chem. 2016;291(10):5247–58.

5. Knupp C, Squire JM. A new twist in the collagen story--the type VI segmented supercoil. EMBO J. 2001;20(3):372–6.

6. Beecher N, Roseman AM, Jowitt TA, Berry R, Troilo H, Kammerer RA, et al. Collagen VI, conformation of A-domain arrays and microfibril architecture. J Biol Chem. 2011;286(46):40266–75.

7. Heumuller SE, Talantikite M, Napoli M, Armengaud J, Morgelin M, Hartmann U, et al. C-terminal proteolysis of the collagen VI alpha3 chain by BMP-1 and proprotein convertase(s) releases endotrophin in fragments of different sizes. J Biol Chem. 2019;294(37):13769–80.

8. Park J, Scherer PE. Adipocyte-derived endotrophin promotes malignant tumor progression. J Clin Invest. 2012;122(11):4243–56.

9. Sun K, Park J, Gupta OT, Holland WL, Auerbach P, Zhang N, et al. Endotrophin triggers adipose tissue fibrosis and metabolic dysfunction. Nat Commun. 2014;5:3485.

10. Li X, Zhao YS, Chen C, Yang L, Lee HH, Wang ZN, et al. Critical Role of Matrix Metalloproteinase 14 in Adipose Tissue Remodeling during Obesity. Molecular and Cellular Biology. 2020;40(8).

11. Bonnemann CG. The collagen VI-related myopathies: muscle meets its matrix. Nat Rev Neurol. 2011;7(7):379–90.

12. Scacheri PC, Gillanders EM, Subramony SH, Vedanarayanan V, Crowe CA, Thakore N, et al. Novel mutations in collagen VI genes: expansion of the Bethlem myopathy phenotype. Neurology. 2002;58(4):593–602.

13. Merlini L, Martoni E, Grumati P, Sabatelli P, Squarzoni S, Urciuolo A, et al. Autosomal recessive myosclerosis myopathy is a collagen VI disorder. Neurology. 2008;71(16):1245–53.

14. Zech M, Lam DD, Francescatto L, Schormair B, Salminen AV, Jochim A, et al. Recessive mutations in the alpha3 (VI) collagen gene COL6A3 cause early-onset isolated dystonia. Am J Hum Genet. 2015;96(6):883–93.

15. Irwin WA, Bergamin N, Sabatelli P, Reggiani C, Megighian A, Merlini L, et al. Mitochondrial dysfunction and apoptosis in myopathic mice with collagen VI deficiency. Nat Genet. 2003;35(4):367–71.

16. Grumati P, Coletto L, Sabatelli P, Cescon M, Angelin A, Bertaggia E, et al. Autophagy is defective in collagen VI muscular dystrophies, and its reactivation rescues myofiber degeneration. Nat Med. 2010;16(11):1313–20.

17. Pfaff M, Aumailley M, Specks U, Knolle J, Zerwes HG, Timpl R. Integrin and Arg-Gly-Asp dependence of cell adhesion to the native and unfolded triple helix of collagen type VI. Exp Cell Res. 1993;206(1):167–76.

18. Nishiyama A, Stallcup WB. Expression of NG2 proteoglycan causes retention of type VI collagen on the cell surface. Mol Biol Cell. 1993;4(11):1097–108.

19. Nanda A, Carson-Walter EB, Seaman S, Barber TD, Stampfl J, Singh S, et al. TEM8 interacts with the cleaved C5 domain of collagen alpha 3(VI). Cancer Res. 2004;64(3):817–20.

20. Burgi J, Kunz B, Abrami L, Deuquet J, Piersigilli A, Scholl-Burgi S, et al. CMG2/ANTXR2 regulates extracellular collagen VI which accumulates in hyaline fibromatosis syndrome. Nat Commun. 2017;8:15861.

21. Stranecky V, Hoischen A, Hartmannova H, Zaki MS, Chaudhary A, Zudaire E, et al. Mutations in ANTXR1 Cause GAPO Syndrome. American Journal of Human Genetics. 2013;92(5):792–9.

22. Lee JY, Tsai YM, Chao SC, Tu YF. Capillary morphogenesis gene-2 mutation in infantile systemic hyalinosis: ultrastructural study and mutation analysis in a Taiwanese infant. Clin Exp Dermatol. 2005;30(2):176–9.

23. Glover MT, Lake BD, Atherton DJ. Infantile systemic hyalinosis: newly recognized disorder of collagen? Pediatrics. 1991;87(2):228–34.

24. Besschetnova TY, Ichimura T, Katebi N, St Croix B, Bonventre JV, Olsen BR. Regulatory mechanisms of anthrax toxin receptor 1-dependent vascular and connective tissue homeostasis. Matrix Biology. 2015;42:56–73.

25. Nielsen SH, Edsfeldt A, Tengryd C, Gustafsson H, Shore AC, Natali A, et al. The novel collagen matrikine, endotrophin, is associated with mortality and cardiovascular events in patients with atherosclerosis. J Intern Med. 2021;290(1):179–89.

26. Bell SE, Mavila A, Salazar R, Bayless KJ, Kanagala S, Maxwell SA, et al. Differential gene expression during capillary morphogenesis in 3D collagen matrices: regulated expression of genes involved in basement membrane matrix assembly, cell cycle progression, cellular differentiation and G-protein signaling. J Cell Sci. 2001;114(Pt 15):2755–73.

27. Sergeeva OA, van der Goot FG. Converging physiological roles of the anthrax toxin receptors. F1000Res. 2019;8.

28. Bradley KA, Mogridge J, Mourez M, Collier RJ, Young JA. Identification of the cellular receptor for anthrax toxin. Nature. 2001;414(6860):225–9.

29. Liu S, Crown D, Miller-Randolph S, Moayeri M, Wang H, Hu H, et al. Capillary morphogenesis protein-2 is the major receptor mediating lethality of anthrax toxin in vivo. Proceedings of the National Academy of Sciences. 2009;106(30):12424–9.

30. Schiavinato A, Przyklenk M, Kobbe B, Paulsson M, Wagener R. Collagen type VI is the antigen recognized by the ER-TR7 antibody. Eur J Immunol. 2021.

31. Stallcup WB, Dahlin K, Healy P. Interaction of the NG2 chondroitin sulfate proteoglycan with type VI collagen. Journal of Cell Biology. 1990;111(6):3177–88.

32. Aigner T, Hambach L, Söder S, Schlötzer-Schrehardt U, Pöschl E. The C5 Domain of Col6A3 Is Cleaved Off from the Col6 Fibrils Immediately after Secretion. Biochemical and Biophysical Research Communications. 2002;290(2):743–8.

33. Tillet E, Ruggiero F, Nishiyama A, Stallcup WB. The Membrane-spanning Proteoglycan NG2 Binds to Collagens V and VI through the Central Nonglobular Domain of Its Core Protein. Journal of Biological Chemistry. 1997;272(16):10769–76.

34. Burg MA, Tillet E, Timpl R, Stallcup WB. Binding of the NG2 Proteoglycan to Type VI Collagen and Other Extracellular Matrix Molecules. Journal of Biological Chemistry. 1996;271(42):26110–6.

35. Mayer U, Poschl E, Nischt R, Specks U, Pan TC, Chu ML, et al. Recombinant expression and properties of the Kunitz-type protease-inhibitor module from human type VI collagen alpha 3(VI) chain. Eur J Biochem. 1994;225(2):573–80.

36. Williams L, Layton T, Yang N, Feldmann M, Nanchahal J. Collagen VI as a driver and disease biomarker in human fibrosis. Febs Journal. 2021.

37. Scherer PE, Williams S, Fogliano M, Baldini G, Lodish HF. A Novel Serum-Protein Similar to C1q, Produced Exclusively in Adipocytes. Journal of Biological Chemistry. 1995;270(45):26746–9.

38. Maeda N, Funahashi T, Matsuzawa Y, Shimomura I. Adiponectin, a unique adipocyte-derived factor beyond hormones. Atherosclerosis. 2020;292:1–9.

39. Yamauchi T, Kamon J, Ito Y, Tsuchida A, Yokomizo T, Kita S, et al. Cloning of adiponectin receptors that mediate antidiabetic metabolic effects. Nature. 2003;423(6941):762–9.

40. Hug C, Wang J, Ahmad NS, Bogan JS, Tsao TS, Lodish HF. T-cadherin is a receptor for hexameric and high-molecular-weight forms of Acrp30/adiponectin. Proceedings of the National Academy of Sciences of the United States of America. 2004;101(28):10308–13.

41. Zweckstetter M, Czisch M, Mayer U, Chu M-L, Zinth W, Timpl R, et al. Structure and multiple conformations of the Kunitz-type domain from human type VI collagen α3(VI) chain in solution. Structure. 1996;4(2):195–209.

42. Bu DW, Crewe C, Kusminski CM, Gordillo R, Ghaben AL, Kim M, et al. Human endotrophin as a driver of malignant tumor growth. Jci Insight. 2019;4(9).

43. Wishart AL, Conner SJ, Guarin JR, Fatherree JP, Peng YF, McGinn RA, et al. Decellularized extracellular matrix scaffolds identify full-length collagen VI as a driver of breast cancer cell invasion in obesity and metastasis. Sci Adv. 2020;6(43).

44. Reeves CV, Wang X, Charles-Horvath PC, Vink JY, Borisenko VY, Young JAT, et al. Anthrax Toxin Receptor 2 Functions in ECM Homeostasis of the Murine Reproductive Tract and Promotes MMP Activity. PLoS ONE. 2012;7(4):e34862.

45. Oh J, Kim CS, Kim M, Jo W, Sung YH, Park J. Type VI collagen and its cleavage product, endotrophin, cooperatively regulate the adipogenic and lipolytic capacity of adipocytes. Metabolism. 2021;114.

46. Lamande SR, Morgelin M, Adams NE, Selan C, Allen JM. The C5 domain of the collagen VI alpha3(VI) chain is critical for extracellular microfibril formation and is present in the extracellular matrix of cultured cells. J Biol Chem. 2006;281(24):16607–14.

47. Maertens B, Hopkins D, Franzke CW, Keene DR, Bruckner-Tuderman L, Greenspan DS, et al. Cleavage and oligomerization of gliomedin, a transmembrane collagen required for node of ranvier formation. J Biol Chem. 2007;282(14):10647–59.

48. Moya-Torres A, Gupta M, Heide F, Krahn N, Legare S, Nikodemus D, et al. Homogenous overexpression of the extracellular matrix protein Netrin-1 in a hollow fiber bioreactor. Applied Microbiology and Biotechnology. 2021.

49. Yuan P, Zhang H, Cai C, Zhu S, Zhou Y, Yang X, et al. Chondroitin sulfate proteoglycan 4 functions as the cellular receptor for Clostridium difficile toxin B. Cell Research. 2015;25(2):157–68.

50. Sengle G, Charbonneau NL, Ono RN, Sasaki T, Alvarez J, Keene DR, et al. Targeting of Bone Morphogenetic Protein Growth Factor Complexes to Fibrillin. Journal of Biological Chemistry. 2008;283(20):13874–88.

